# Multi-omic data fusion reveals the *in vivo* enzyme kinetics of *Vibrio natriegens* at the genome-scale

**DOI:** 10.64898/2026.06.02.729587

**Authors:** St. Elmo Wilken, Martin Beyß, Miroslav Kratochvíl, Alexa Barbara Grebel, Karen Methling, Anja Stefanski, Paul London, Michael Lalk, Klaus Schaper, Ilka Maria Axmann, Katharina Nöh, Philipp Westhoff, Oliver Ebenhöh

## Abstract

*Vibrio natriegens* is a halophilic, Gram-negative marine bacterium that is increasingly used in metabolic engineering applications due to its fast growth rate. In sparse minimal medium the organism has a doubling time of 25 minutes, which is about twice as fast as *Escherichia coli* under similar conditions. Given that its protein density is similarly constrained to that of *E. coli*, this necessitates that its metabolic enzymes are able to catalyze flux at a higher rate to sustain its metabolism. In this work, we measure the apparent turnover numbers of metabolically active enzymes in *V. natriegens* under a variety of growth conditions.

The apparent turnover numbers of *V. natriegens* enzymes were measured *in vivo* by conducting coupled quantitative proteomics and ^13^C metabolic flux analysis experiments under seven different carbon source conditions in sparse minimal medium. A high quality genome-scale metabolic model was constructed and curated using additional experimental data. This model was extended with enzyme constraints, and subsequently used to find kinetic parameters that minimize the difference between model predictions and experimental observations. This model guided data fusion approach enabled the estimation of 357 apparent turnover numbers for metabolically active enzymes in *V. natriegens*.

Our results reveal that the metabolic enzymes of *V. natriegens* are in median 14-fold faster than those of *E. coli* under similar conditions. Moreover, we show that machine learning generated turnover number estimates substantially underestimate the kinetics of *V. natriegens*. Our turnover number estimates were used to parameterize multiple condition dependent enzyme constrained flux balance analysis models of *V. natriegens*, which improved their predictive accuracy compared to the machine learning parameterisation. The combined experimental-computational approach employed here sheds light on the mechanism *V. natriegens* uses to accelerate its growth. This approach can also be extended to other bacteria, increasing the availability of *in vivo* measured enzyme turnover numbers, and improving the predictive accuracy of enzyme constrained metabolic models of other microbes.

## 1 Introduction

*Vibrio natriegens* is a halophilic, Gram-negative marine bacterium, well known for its fast growth rate. Under optimal conditions its doubling time is less than 10 minutes in complex medium [1]. Even when cultured in a sparse minimal medium the organism reaches doubling times of 25 minutes, which is competitive with *Escherichia coli* in complex medium [2]. This raises the question: what adaptations does *V. natriegens* possess that allow it to grow so fast?

Protein synthesis is a major biosynthetic cost for rapidly growing cells [3], necessitating the regulation of high fluxes towards both ATP production and amino acids to supply ribosomes with sufficient precursors to polymerize proteins. Catabolic and anabolic enzymes are used to drive these processes, but their expression is regulated according to the growth rate of the organism [4]. Detailed measurements have revealed that the cellular density of bacteria is relatively constant [5], leading to the emergence of a trade-off between devoting limited space to metabolic enzymes or ribosomes. Fundamentally, the growth rate of an organism is bounded by the rate at which its ribosomes can synthesize new proteins (including ribosomal proteins). At optimal, balanced growth the flux of proteinogenic precursors must match their consumption rate by ribosomes. At higher growth rates more ribosomes are needed, but then less space remains for metabolic enzymes. As a consequence, faster growth is limited to richer nutrient conditions that enable cells to invest more resources into ribosomes, and less into metabolic enzymes [6].

Assuming that a growth rate independent total protein capacity limitation also exists in *V. natriegens*, the organism must increase the kinetic speed of its enzymes to accommodate for the biosynthetic cost of its substantially faster growth rate compared to *E. coli* in comparable conditions. Recent measurements have revealed that the ribosomal translation rate of *V. natriegens* is approximately 40% faster than that of *E. coli* [7]. While the metabolic enzyme kinetics were not measured, it is implied that they too must be faster to explain the enhanced growth rate of *V. natriegens* [7].

Enzyme kinetics can be described by Michaelis-Menten like rate laws, where the enzyme turnover number, k_cat_, represents the maximal reaction rate an enzyme can achieve [8]. The turnover number can be measured *in vitro*, but this approach is laborious (e.g. it requires careful buffer optimization for each measurement), and is unsuitable for high throughput analyses. An alternative method to estimate enzyme turnover numbers uses ^13^C metabolic flux analysis (^13^C MFA) to measure the intracellular flux, *𝓋*, catalyzed by an enzyme, and quantitative proteomics to measure the concomitant amount of the enzyme, *e*, required to sustain this flux *in vivo* [9, 10]. *The apparent turnover number, k*_*app*_, *is defined as the ratio of the flux and enzyme amount required for a reaction [11], and lumps the true turnover number together with saturation, regulation, and thermodynamic effects, η, as shown in Equation (1)*.

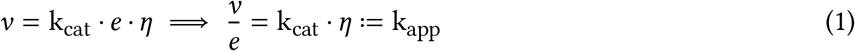

In theory, the maximum observed apparent turnover number (k_max_) across a wide range of conditions should approximate the true underlying enzyme turnover number [9]. An advantage of measuring in vivo apparent turnover numbers is that physiological effects are naturally incorporated into the measurement cf. in vitro data which lack cellular context, and often rely on heterologously expressed enzymes that may be missing endogenous post-translational modifications.

In this work, we combine *in vivo* intracellular flux measurements, derived from ^13^C MFA, and quantitative proteomics of *V. natriegens* across seven carefully chosen media conditions to probe its metabolism. To expand the scope of the intracellular flux measurements beyond the core central carbon metabolism, we also reconstructed a new, detailed genome-scale metabolic model of *V. natriegens*. This model was extended with enzyme constraints, and an efficient model-based parameter optimization scheme was developed that fuses the proteomic and fluxomic data to find kinetic parameters that minimize the discrepancy between model predictions and experimental observations at both the fluxome and proteome levels [12]. This enabled us to estimate 357 *in vivo* apparent turnover numbers spanning the primary metabolism of *V. natriegens*. Comparing these data to similar estimates for *E. coli*, we found that *V. natriegens* has enzymes that are kinetically 14-fold faster than those of *E. coli* in median, partially explaining its enhanced growth rate.

The apparent turnover numbers found here substantially deviated from machine learning-based predictions using enzyme sequence and reaction participant data. Parameterizing a condition dependent enzyme constrained flux balance analysis model with the apparent turnover numbers found here significantly improved the predictive performance of the model compared to using only the machine learning based estimates. The approach applied here can be easily extended to other organisms and highlights the increasing importance of fusing heterogenous omics data to elucidate complex metabolic phenomena and improve the predictive accuracy of metabolic models.

## 2 Results

Figure 1 presents an overview of the methodology developed in this work to estimate the enzyme kinetic parameters of *V. natriegens*. First, quantitative proteomic data are used to measure the proteome allocation of *V. natriegens* under seven different growth conditions. Second, metabolomic and absolute fluxomic data quantify the activity of its metabolic enzymes under each of these conditions. Thereafter, these data are combined with an enzyme constrained genome-scale metabolic model that is parameterized by a set of enzyme kinetic constants (k_app_ values). An efficient gradient-based optimization schemes updates these k_app_ values to minimize the loss between model predictions and experimental observations. The predictive value of the k_app_ estimates is investigated. Finally, comparing the k_app_ estimates to machine learning based k_cat_ predictions underscores the importance of using multi-omic data fusion for parameter estimation of biological models.

**Figure 1:**
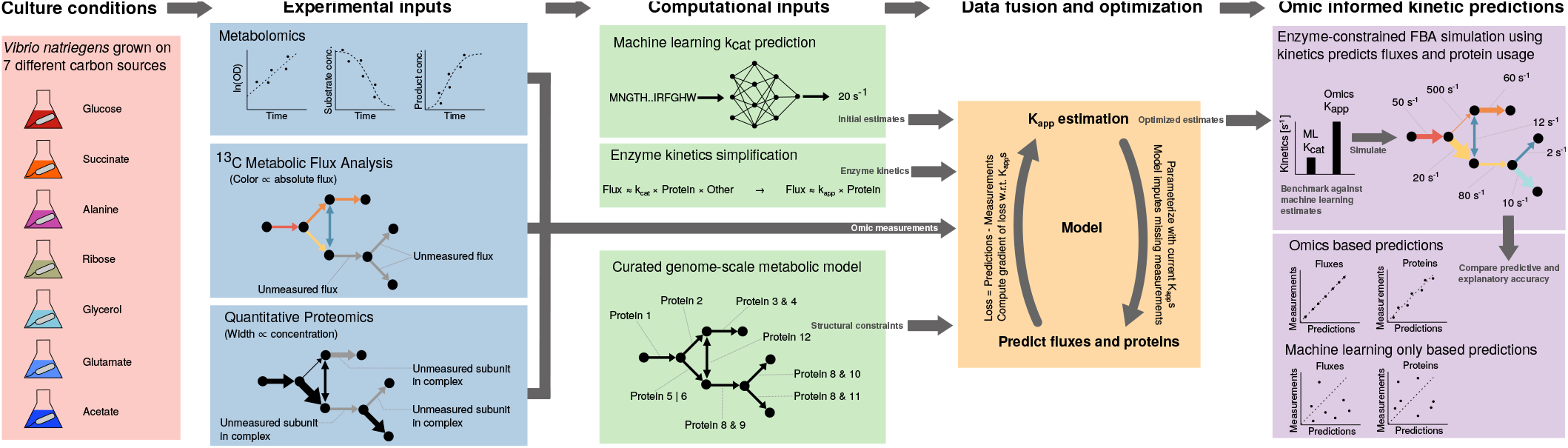
Multi-omic data fusion integrates a genome-scale metabolic model with high-throughput measurements for parameter estimation. An overview of the methodology developed in this work to estimate enzyme kinetics using omics data is presented here. *V. natriegens* was grown in seven different media conditions, in which metabolomic, fluxomic, and proteomic data were measured. A curated enzyme constrained genome-scale metabolic model was reconstructed, and used to find parameters that minimize the discrepancy (loss) between model predictions and experimental observations. The optimization scheme is iterative, and makes use of efficiently calculated gradients to rapidly minimize the loss. Machine learning generated kinetics were used as initial estimates for the parameter optimization scheme. The optimal set of kinetics were found to have significantly better explanatory and predictive validity than the machine learning generated estimates.

### Protein capacity limitations leads to resource allocation constraints

Cellular growth rate is an important regulator of gene expression in bacteria. Fast growing bacteria have evolved global regulatory mechanisms to adapt their proteome under different nutrient conditions to avoid cellular crowding. The protein density of *E. coli* has been found to vary by less than 10 % across a range of media conditions and growth rates, which leads to a resource allocation trade-off [13]. Since translation rate does not vary strongly with growth rate [14], faster growth necessitates more resources to be devoted to ribosomes for increased protein synthesis. To investigate whether *V. natriegens* is also subject to a total protein capacity bound, we measured its protein mass fraction across seven conditions in defined, minimal VN medium [2]. For each carbon source, we achieved growth rates comparable to previously published data [2]: glucose (1.74 ± 0.04 h^−1^), succinate (1.05 ± 0.05 h^−1^), alanine (0.95 ± 0.02 h^−1^), ribose (0.87 ± 0.03 h^−1^), glycerol (0.77 ± 0.01 h^−1^), glutamate (0.67 ± 0.02 h^−1^), and acetate (0.63 ± 0.01 h^−1^). Similarly to *E. coli*, the total protein density of *V. natriegens* is independent of growth rate (Figure S1), supporting the observation that protein capacity limitations are a universal bacterial phenomenon. We find that the mean protein density of *V. natriegens* is slightly less than that of *E. coli* (0.46 ± 0.03 g g_DW_^−1^ cf. 0.55 g g_DW_^−1^ respectively), but in agreement with a previous *V. natriegens* estimate in glucose minimal media [15, 16].

Given this reduced proteome capacity constraint, we next investigated how *V. natriegens* allocates its proteome to achieve high growth rates in minimal medium. We conducted quantitative proteomics experiments using the same seven media conditions, harvesting cells in mid-exponential phase (see Supplementary Section S2.1 for more details, and Supplementary Data 1 for the quantitative proteome). A hallmark consequence of the total proteome capacity limitation is a linear correlation between ribosome mass fraction and growth rate [17]. Figure 2A shows that *V. natriegens* exhibits a linear relationship between growth rate and translational proteins (mostly composed of ribosomal proteins), as has been found in many other bacteria [18]. At the maximum growth rate investigated in this work (carbon source: glucose), approximately 27 % of the total proteome budget is allocated to translation machinery. These findings are consistent with a previous dataset for *V. natriegens* [7].

**Figure 2:**
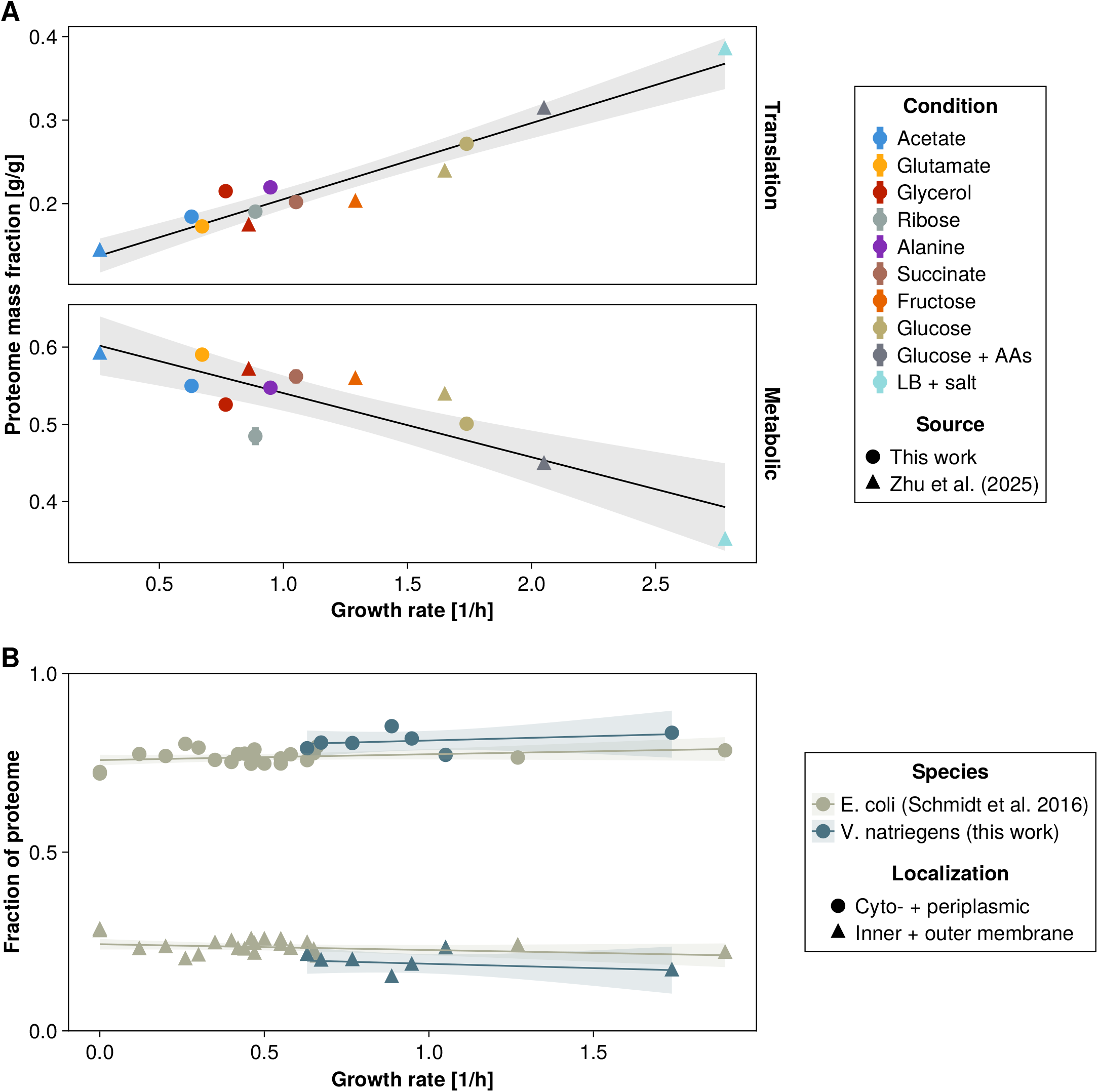
Proteome allocation of *V. natriegens* is strongly coupled to growth rate. The trade-off between ribosomal and metabolic proteome mass fractions follow the same trends as seen in *E. coli*, including the partitioning between cytosolic and membrane proteins. **A:** Higher growth rates demand more ribosomal investment, reducing the space available for other metabolic proteins. Proteome mass fraction allocated to translation associated proteins, based on COG classification (category J), for *V. natriegens* under seven different media conditions. Regression slope statistically significant. Here, metabolic enzymes are defined as all proteins assigned to the COG categories: carbohydrate (G), amino acid (E), nucleotide (F), lipid (I), inorganic ion (P), energy (C), and coenzyme (H) metabolism. **B:** In contrast to metabolic proteins, there is no statistically significant trend between the proteins assigned to the cytoplasm and periplasm, or membrane compartments as growth rate changes. Localization of each protein was predicted with DeepLocPro [21]. Each point represents the mean of at least 3 biological replicates, and the error bar presents 1 standard deviation (note, the errors are smaller than the markers in the figure, and thus not visible). Trend lines are linear regressions, with the shaded region indicating the 95% confidence intervals. Regression slope significance tests were conducted using the Wald test, with a p-value threshold of 0.05. Each data point is the sum of all proteins assigned to the respective category. The carbon source order in the legend is with respect to increasing growth rate. Zhu et al. (2025) and Schmidt et al. (2016) are a previously published proteomic datasets for *V. natriegens* and *E. coli* under similar conditions, respectively [7, 22].

Next, we investigated the compartmentalization of proteome mass across growth rates. Interestingly, we find that the total membrane associated protein mass fraction for *V. natriegens* stays relatively constant across all seven culture conditions, in agreement with data from *E. coli* (statistically insignificant slopes, Figure 2B). Additionally, we find that *V. natriegens* allocates approximately the same mass fraction to membrane proteins as *E. coli* does (0.19 ± 0.03 g g^−1^ cf. 0.23 ± 0.02 g g^−1^ respectively), which may be attributable to their similar rod shape and size (3 µm^3^ to 4 µm^3^ during fast exponential growth) [19, 20].

### ^13^C MFA reveals sustained high intracellular metabolic fluxes in *V. natriegens*

A metabolic model of *V. natriegens*’s central carbon metabolism was formulated (Supplementary Data 2), covering all central carbon pathways and biomass synthesis. The model was constructed to accommodate all available ^13^C labeling data, including growth on the same seven substrates used in the proteomic measurements: glucose, succinate, alanine, ribose, glycerol, glutamate, and acetate (Supplementary Figure S15). Bioprocess models were used to estimate extracellular fluxes based on time course ^1^H-NMR measurements of the extracellular metabolites (Supplementary Data 3) in each carbon source condition, and to evaluate metabolic stationarity for flux estimation (Supplementary Figures S13 and S14). Careful experimental design ensured that the measurements took place during isotopic steady state thereby fulfilling the requirements for ^13^C MFA (see the materials and method section). ^13^C tracer incorporation into biomass bound amino acids, together with the extracellular fluxes, were incorporated into the ^13^C MFA model, which was used to estimate intracellular metabolic fluxes under the seven different carbon source conditions used in this work. No significant fit for the glycerol condition could be found, hinting that *V. natriegens* might use a non-canonical glycerol metabolism. For all the other six conditions a robust fit was found, and all the subsequent work only used these six conditions. A Bayesian approach was employed to generate flux estimates congruent with the measured data as described in the materials and methods section. See Supplementary Figures S16 – S21 for visualizations of the estimated intracellular fluxes including their uncertainty and Supplementary Figures S22 – S27 for the marginal posterior distributions.

Figure 3 summarizes the estimated intracellular fluxes of *V. natriegens* subject to the same culture conditions used in the proteomic analyses of Figure 2. The fluxes in Figure 3 are normalized to the uptake rate of each respective carbon source, and the error bars denote the 95 % credible intervals based on the marginal posterior flux estimates (reported in Supplementary Data 4). The named reactions are visualized in the network map in Supplementary Figure S28. The credible intervals for most metabolic fluxes are tight, with notable exceptions in the cata/anaplerotic reactions for succinate, glutamate, and acetate driven growth, as well as the pentose phosphate pathway (PPP) for ribose driven growth. In the latter case, the flux intervals are relatively large, but in most cases only the magnitude is uncertain, not the direction.

**Figure 3:**
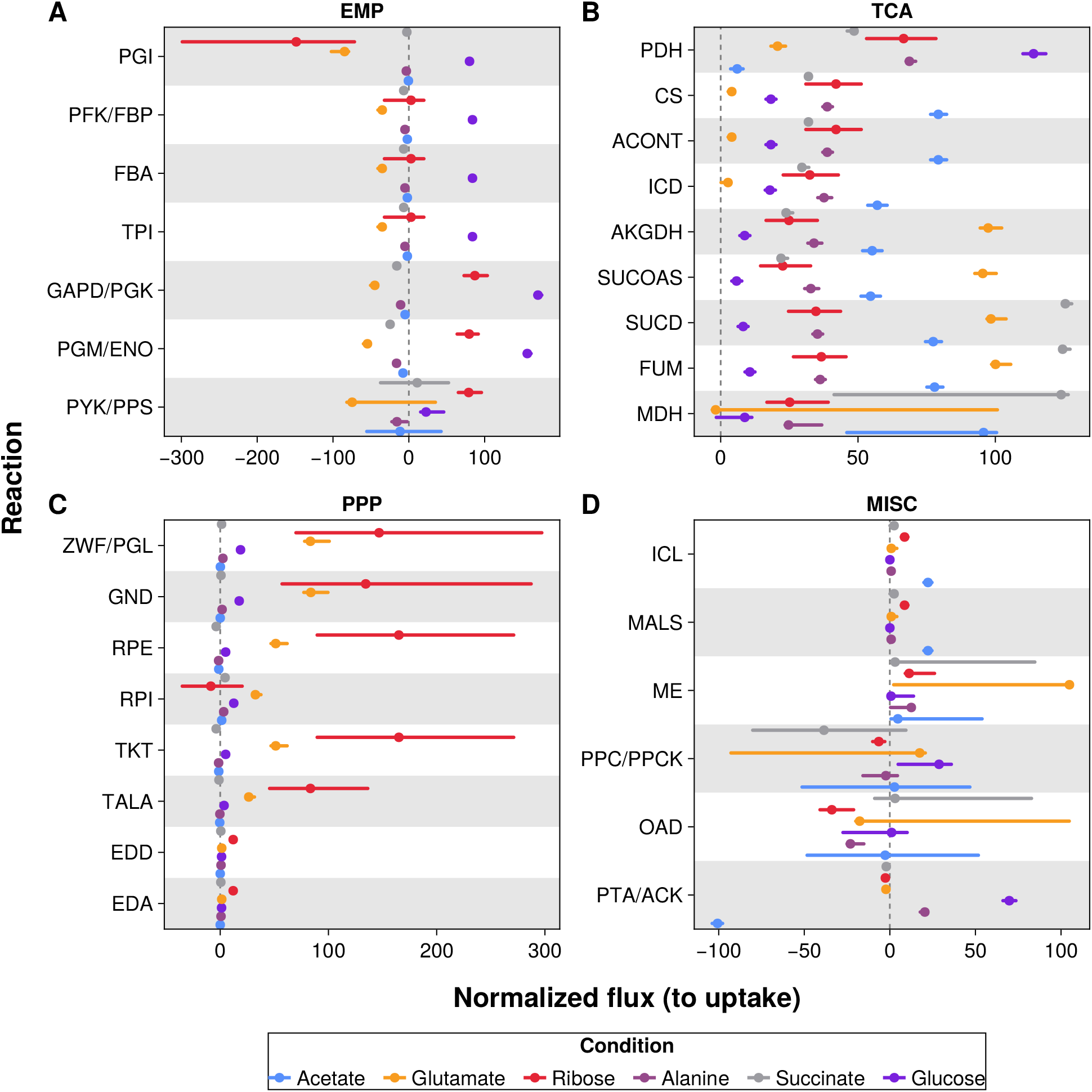
^13^C MFA intracellular flux estimates for *V. natriegens* in six different culture conditions in defined medium. Fluxes are normalized to the uptake rate of the respective carbon source. Mode of the marginal posterior flux estimates are shown as points, and the 95 % credible interval is shown in the error bars. **A:** Glycolysis/Embden-Meyerhof-Parnas grouping (EMP) **B:** Citric acid cycle grouping (TCA) **C:** Pentose phosphate pathway grouping (PPP) **D:** Anaplerotic, glyoxylate shunt, and acetate overflow are grouped as miscellaneous reactions (MISC). The named reactions are visualized in the network map in Supplementary Figure S28. ACONT: Aconitate hydratase, AKGDH: 2-Oxogluterate dehydrogenase, CS: Citrate synthase, EDA: 2-dehydro-3-deoxy-phosphogluconate aldolase, EDD: 6-phosphogluconate dehydratase, FBA: Fructose-bisphosphate aldolase, FUM: Fumarase, MDH: Malate dehydrogenase, GAPD/PGK: Glyceraldehyde-3-phosphate dehydrogenase/Phosphoglycerate kinase GND: Phosphogluconate dehydrogenase, ICD: Isocitrate dehydrogenase, ICL: Isocitrate lyase, MALS: Malate synthase, ME: Malic enzyme, OAD: Oxaloacetate decarboxylase, PDH: Pyruvate dehydrogenase, PFK/FBP: Phosphofructokinase/Fructose-bisphosphatase, PGI: Glucose-6-phosphate isomerase, PGM/ENO: Phosphoglycerate mutase/Enolase, PPC/PPCK: Phosphoenolpyruvate carboxylase/Phosphoenolpyruvate carboxykinase, PTA/ACK: Phosphotransacetylase/Acetate kinase, PYK/PPS: Pyruvate kinase/Phosphoenolpyruvate synthase, RPE: Ribulose 5-phosphate 3-epimerase, RPI: Ribose-5-phosphate isomerase, SUCD: Succinate dehydrogenase, SUCOAS: Succinyl-CoA synthetase, TALA: Transaldolase, TKT: Transketolase, TPI: Triose-phosphate isomerase, ZWF/PGL: Glucose 6-phosphate dehydrogenase/6-phosphogluconolactonase.

Considering growth using glucose as the sole carbon source at first, the intracellular flux estimates reported here of *V. natriegens* match those of a previous study [15](Supplementary Figure S29). Recapitulating previous results, we also find that the metabolism of *V. natriegens* resembles that of wild type *E. coli* MG1655 (growth rate 0.68 h^−1^ in comparable conditions). The most prominent differences are that *E. coli* channels relatively more flux through the PPP than *V. natriegens*, but the relative flux differences are smaller in the citric acid cycle (tricarboxylic acid cycle, TCA) and the Embden-Meyerhof-Parnas pathways (EMP, glycolysis), as shown in Supplementary Figure S30. Interestingly, by further comparing *V. natriegens* to an *E. coli* MG1655 that was evolved for faster growth (growth rate 0.94 h^−1^ in comparable conditions) [23], we see that the discrepancy in relative PPP fluxes reduces, and their metabolism converges more (Supplementary Figure S30). In contrast, intracellular flux data for the faster growing *Geobacillus* LC300 (growth rate 2.15 h^−1^ vs. *V. natriegens* 1.74 h^−1^ in similar conditions) [24] reveals that it channels substantially more relative flux through the PPP, suggesting that flux through the PPP is not a large factor in the growth rate differences between these three organisms. On an absolute basis, faster growth rates are associated with higher fluxes, but when normalizing by growth rate the interspecies flux distribution differences disappear (Supplementary Figures S31 and S32). This suggests that the ability to channel higher fluxes through metabolism, and not unique flux configurations, is a better explanatory variable for the observed growth rate differences between the organisms.

Both glucose and ribose driven growth are glycolytic based on the intracellular flux directions. In contrast, succinate, alanine, glutamate, and acetate represent gluconeogenic substrates, as confirmed by the flux estimates in the EMP pathway (glycolysis) as well as the proteomic data. Growth on these latter substrates exhibit increased fluxes through the TCA cycle where they enter metabolism. Acetate in particular, a purely respirative carbon source, is associated with a significantly upregulated TCA metabolism across the board. Succinate shows an increased flux through succinate dehydrogenase (127 % of uptake flux, SUCD), which couples the substrate to the electron transport chain. Likewise, the highest relative flux through 2-oxogluterate dehydrogenase (AKGDH) is observed under the glutamate condition, which is its entry point into the core metabolism. Similarly, under ribose driven growth, fluxes through the PPP are substantially higher than in the other carbon conditions because this is the entry point into metabolism for the substrate.

Compared to the other carbon sources, glucose driven growth is associated with the highest intracellular fluxes relative to uptake through the EMP pathway in the glycolytic direction. In the gluconeogenic direction, glutamate drives the highest fluxes through the EMP pathway. Particularly for the ribose condition, the Entner–Doudoroff (ED) pathway is the most active (11 % of uptake flux) compared to all the other carbon sources. There is appreciable flux through the glyoxylate shunt (8 % of uptake flux) in this condition, but this is dwarfed by the acetate condition, which channels 22 % of its uptake flux through the shunt to converse carbon and generate acetyl coenzyme A. For the ribose condition, the cata/anaplerotic fluxes are well resolved, revealing significant fluxes relative to ribose uptake: oxaloacetate decarboxylase (30 %), phosphoenolpyruvate carboxykinase (6 %) and malic enzyme (17 %). Interestingly, these reactions potentially create a futile cycle.

The substrate uptake rates are high for the glycolytic substrates (glucose: 21 mmol g_DW_^−1^ h^−1^, ribose: 16 mmol g_DW_^−1^ h^−1^), and the gluconeogenic substrates (acetate: 39 mmol g_DW_^−1^ h^−1^, alanine: 30 mmol g_DW_^−1^ h^−1^, succinate: 23 mmol g_DW_^−1^ h^−1^, glutamate: 15 mmol g_DW_^−1^ h^−1^), which leads to concomitantly high intracellular fluxes, typical of *V. natriegens*. In sum, *V. natriegens* is able to consistently channel high fluxes through its metabolism, regardless of the carbon source it is growing on.

### Using model guided data fusion for kinetic parameter estimation

Enzyme constrained genome-scale metabolic models are an extension of classic flux balance type models [25]. In addition to a metabolic network with flux direction constraints, enzyme constrained models include linearized enzyme kinetics: *𝓋*_*i*_ = *k*_cat,*i*_ ⋅ *e*_*i*_, where *𝓋*_*i*_ is the flux, *e*_*i*_ is the enzyme concentration, and *k*_cat,*i*_ is the enzyme turnover number for the associated reaction *i*. Additionally, possibly multiple protein capacity constraints are included: 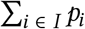⋅ MW_*i*_ ≤ *P*_*I*_ where MW_*i*_ is the molar mass of protein *i* and *I* is some set of proteins that have a bounded mass fraction. Typically, *I* is the set of all proteins in the model, and the bound represents the total metabolic proteome of the cell, i.e. *P*_*I*_ = *P*_total_. In this work, we consider two protein capacity bounds: a total bound (*P*_total_) and a membrane bound (*P*_membrane_) [26]. The latter bound reflects the observation that the fraction of the proteome assigned to the membrane is relatively constant (Figure 2B). Note, enzyme complexes make it necessary to distinguish between proteins, *p*_*i*_ for protein *i*, and enzymes, *e*_*i*_ for enzyme *i*. Multiple (different) proteins may be associated with a single enzyme, but this complexity is abstracted away into the gene reaction rules (GRRs) of the model and captured by the function GRR_*i*_ for enzyme *i*. This function constrains the protein subunit stoichiometry of an enzyme to match the composition of the relevant enzyme in the model. The classic formulation of the enzyme constrained flux balance analysis model maximizing predicted growth rate (*µ*, with *µ* ∈ **v**) is shown in Equation (2). Here **S** represents the stoichiometric matrix of the model (the mole balance equations across each reaction), and **v**_LB_, **v**_UB_ the lower and upper bounds on fluxes respectively.

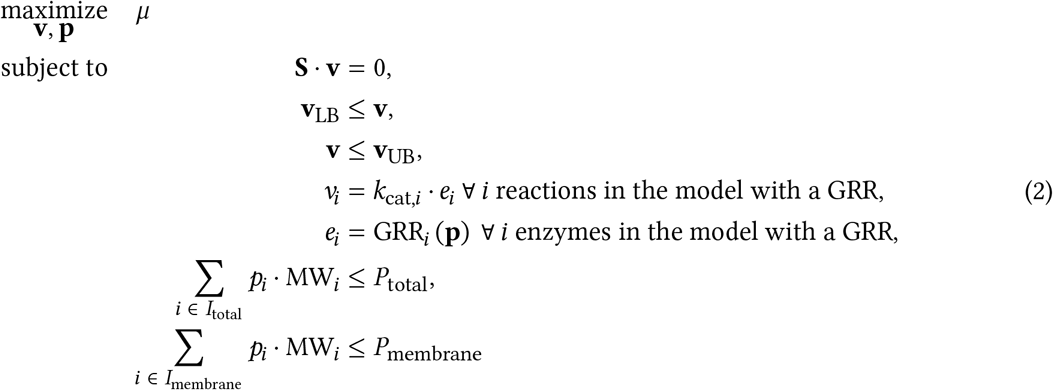

While enzyme constrained models enable better metabolic predictions [27], they require additional parameters that are often challenging to obtain. In particular, the enzyme turnover numbers (k_cat_’s) are laborious to experimentally measure. Recent efforts have focussed on using machine learning models to estimate these constants [28]. In contrast, here we combine quantitative proteomics and ^13^C MFA inferred absolute fluxes with a curated genome-scale metabolic model of *V. natriegens* to estimate these parameters.

First, we define the symbols used in the parameter estimation approach. Let the measured protein abundances be 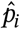 for protein *i* in the set of measurements *M*. Note, given the inherent noisiness associated with proteomic measurements, we included in set *M* only the most abundant proteins measured in each isozyme included in the model. Let the measured (technically inferred from ^13^C MFA) flux, be 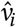 for flux *i* in the set of measured fluxes *F*. This set *F* of measured fluxes is stochastically drawn from the set of Markov Chain Monte Carlo samples comprising the posterior flux distribution (see the material and methods section for more details of the flux estimation procedure). Let 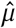 be the measured growth rate.

Second, we simplify the model to enable robust fitting. In general, each reaction may be catalyzed by multiple isozymes, as reflected in the gene reaction rules of the full model, with their own specific kinetics. Here, we simplify this facet of the model, and assume that each isozyme of a specific reaction shares a single kinetic parameter, called the apparent enzyme turnover number *k*_app,*i*_, and that it catalyzes the reaction flux according to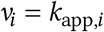 ⋅ *e*_*i*_ for reaction *i*. Note, multiple proteins may be involved in reaction *i*, e.g. in the case of reaction complexes, and this information is captured by the GRRs in the model. For enzyme complexes, only the highest abundance subunit is included in the loss functions, as described below.

Let *K* be the set of reactions for which we will estimate their kinetic parameters using the model and measured data. Suppose an initial estimate of these apparent enzyme turnover numbers (**k**_app_) is given, and these are used to predict the fluxes, **v**, and protein concentrations, **p**, using an enzyme constrained model like Equation (2). The discrepancy between predictions and measurements can be quantified: predicted growth loss through Equation 3, flux loss through Equation 4, measured protein loss through Equation 5. For brevity, define **x** (**k**_app_) = [**v** (**k**_app_) **p** (**k**_app_)], and notice that both the flux and protein predictions depend on **k**_app_, the kinetics of the model.

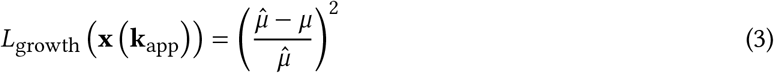

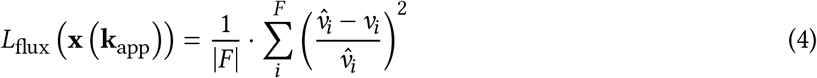

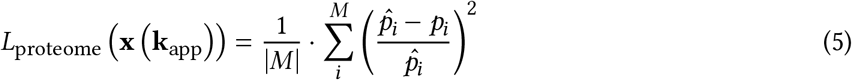

Since we are interested in finding enzyme kinetic parameters that best describe the measured data, we use the total loss, as shown in Equation 6, in the objective of the enzyme constrained model instead of the growth rate. The parameter estimation problem is shown in Equation (7).

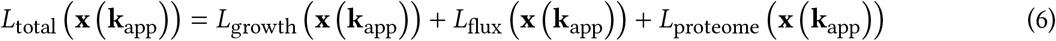

The parameter estimation problem is a bi-level optimization problem. The inner problem of Equation 7 is an enzyme constrained model that predicts the variables, **x**, that minimizes the discrepancy between predictions and experimental measurements given a set of parameters, **k**_app_. In turn, the outer problem seeks to adjust these parameters to yield the best fit to the measured data. Since the inner problem can be efficiently differentiated [12], it is possible to use a first order optimizer to rapidly find kinetic parameters, **k**_app_, that minimizes the discrepancy between model predictions and experimental measurements in the outer problem.

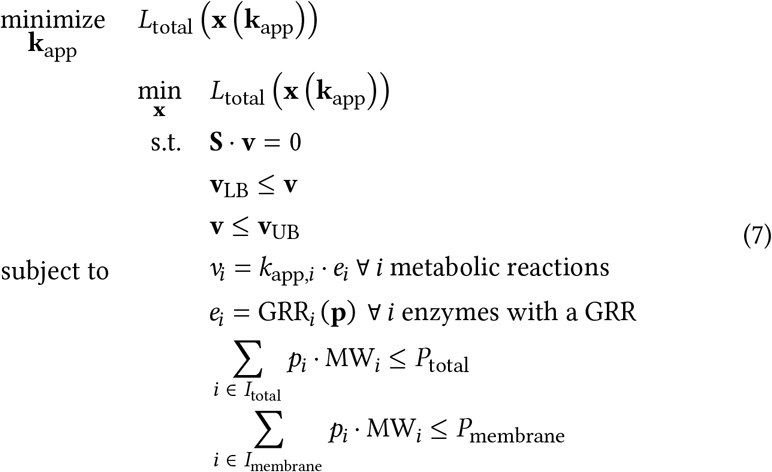

Since a solution to Equation (7) might not be a global minimizer, a multi-start strategy was employed that uses different initial guesses for **k**_app_, based on machine learning estimates of the proteins of *V. natriegens* [29]. Figure 4 shows the five best (lowest total loss at the termination of an optimization run) replicates, where the fluxomic and proteomic losses have been separated for the benefit of the reader, across optimization iterations. It is apparent that the optimization scheme first finds coarse parameters that minimize the flux loss, and then fine tunes them to reduce the protein loss. In all cases the predictive error (loss) is reduced by at least two orders of magnitude from the initial parameterization, suggesting that the model finds k_app_ estimates that explain the observations in the context of Equation (7) well.

**Figure 4:**
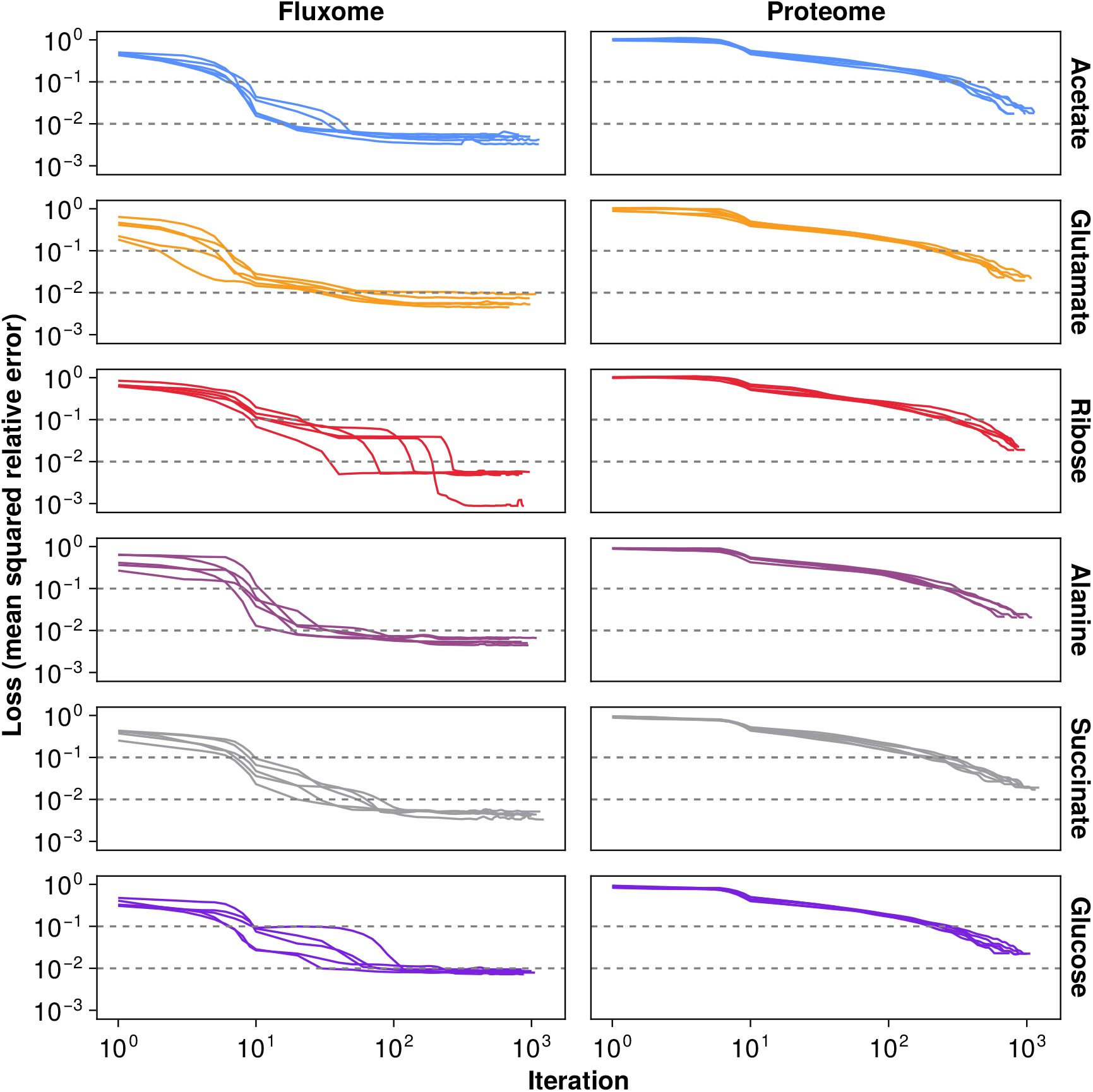
Model parameters are optimized to fit the measured fluxome and proteome for each condition. The fluxome loss represents the mean squared relative error of model predictions relative to the ^13^C MFA estimated fluxes (Equation 4). The proteome loss represents the mean squared relative error of model predictions to the quantitative proteomics data (Equation 5). Only the five best replicates of 200 simulations are shown. Each replicate was initialized with a different random set of parameters. In all conditions, the loss was reduced by at least two orders of magnitude. The optimization termination criterion (absolute change in objective function ≤ 10^−8^) was reached in all five replicates shown.

During the course of the parameter optimization, the predicted variables (fluxes and proteins) rapidly approach their associated measurements for each of the five replicates (Figure 5). For brevity, we initially focus on the glucose condition only, and therein only on the glycolytic pathway, but all the findings extend to the other conditions and pathways as well. Figures 5A and 5B show that the flux and protein predictions narrowly track their measurements despite the initial kinetic parameter estimates yielding high loss predictions. The optimal solutions of Equation (7) yielded mean relative errors for the flux and protein predictions of, respectively, 3 % and 6 % across all the conditions. This is expected as the total loss function includes Equation (4) and Equation (5), which are driven to zero as shown in Figure 4. Figures 5C and 5D extend this analysis to all the measured fluxes and proteins across all the conditions. Here, perfect prediction accuracy should yield data points on the diagonals of the facet plots. The flux estimates track their measurements very closely across all the conditions, with an overall mean relative error of less than 5 %. In contrast, the protein predictions show worse performance with a mean relative error of approximately 211 %. Inspecting the results of Figure 5D closer, it is evident that this large error is attributable to a sizable minority of predictions that overestimate the measurement, and a small minority of predictions that underestimate the measurement (respectively, 9 % and 2 % of predictions are at least a factor of 2 bigger or smaller than the corresponding measurement). These high error proteins belong to enzymes that are heteromeric complexes. The offset is due to the formulation of the protein loss function, where only the highest abundance protein of a complex is included in the loss function, and the subunit stoichiometry is used to determine the concentration of the other subunits. This design choice reflects a compromise to smoothen the objective loss over optimization iterations given the noisiness of proteomics measurements. Excluding these proteins, the overall mean relative error is less than 4 %, which is reflected by the majority of the datapoints falling on the diagonal of Figure 5D. Due to this modest overestimation artifact, the k_app_ estimates derived through the parameter optimization scheme used here likely yield slight underestimates of the true kinetic parameters.

**Figure 5:**
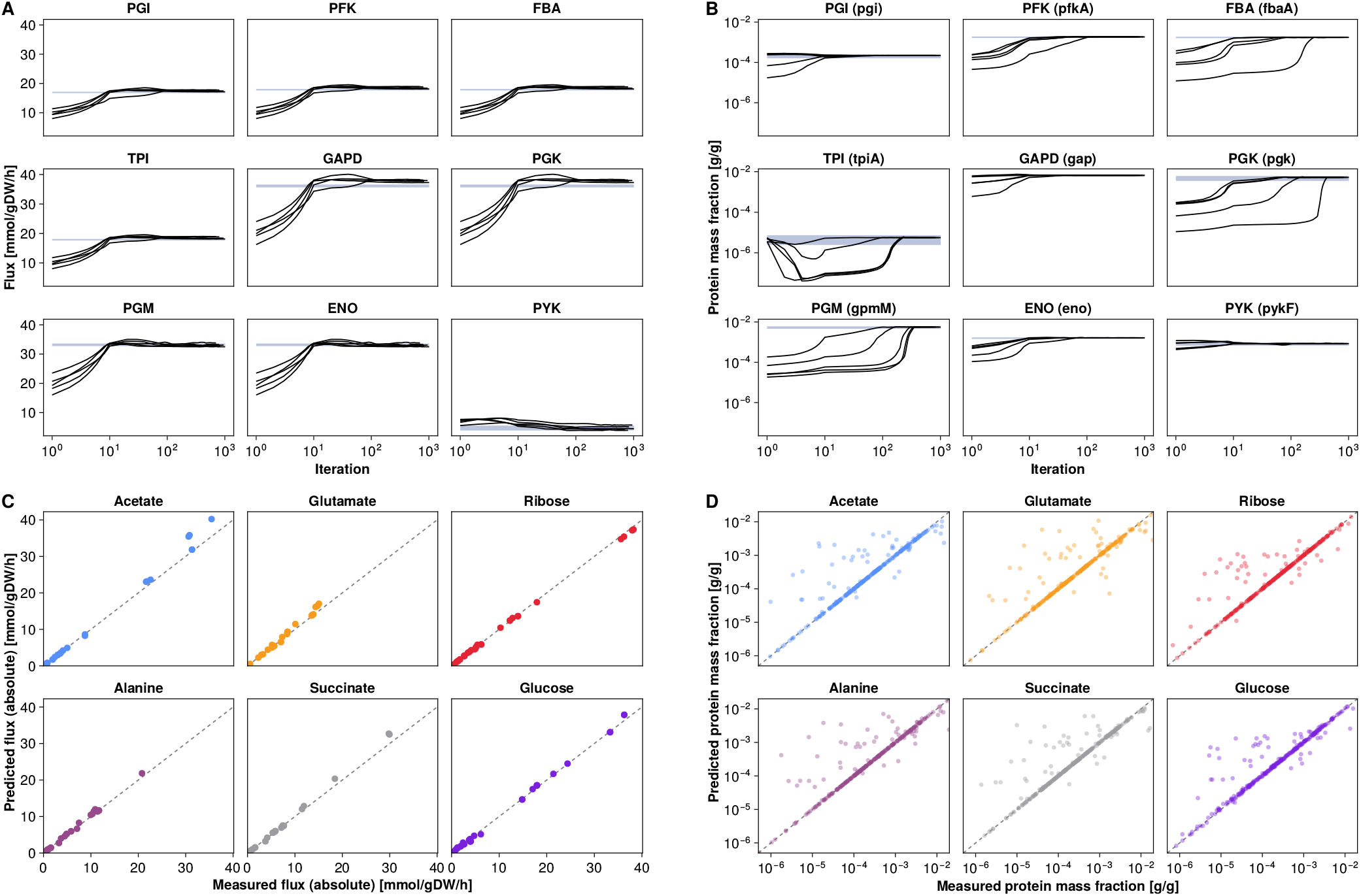
Parameter estimation reduces the discrepancy between model predictions and experimental observations. Each fit of Equation 7 was initialized with a different initial parameterization. The five best (lowest loss for each simulation) fit trajectories are shown in **A** and **B**, while the mean of the five best fit solutions (at the termination of the optimization problem) of the replicates are shown in **C** and **D**. For clarity in **A** and **B**, only the core glycolysis enzymes are shown for the glucose condition, but similar results hold for the other conditions, fluxes, and proteins. **A:** Flux prediction (black line) rapidly converge on the experimentally measured flux (purple shaded region). **B:** Protein concentration predictions also rapidly converge on the experimental measurements (black lines and purple shaded area retain their meaning). For **A** and **B** the reaction is shown in each subplot title, with the associated protein in brackets. **C:** All the measured and predicted fluxes for each condition are shown. **D:** All the measured and predicted protein concentrations for each condition are shown. Due to the structure of the fitting procedure, the fit ends up overestimating the abundance of many proteins, effectively underestimating the final k_app_ estimates. In **C** and **D** the dashed diagonal line represents equality between measurement and prediction. ENO: Enolase, FBA: Fructose-bisphosphate aldolase, GAPD: Glyceraldehyde-3-phosphate dehydrogenase, PGK: Phosphoglycerate kinase, PFK: Phosphofructokinase, PGI: Glucose-6-phosphate isomerase, PGM: Phosphoglycerate mutase, PYK: Pyruvate kinase, TPI: Triose-phosphate isomerase.

While the simulation predictions converge on the measurements (Figure 5), it remains to be seen if the kinetic estimates, the k_app_ values for each reaction, also converge across the replicates. Since Equation 7 is nonlinear, multiple parameter optima may exist given the random initialization that was used. In Figure 6 the convergence of the k_app_ estimates for each replicate across all reactions in the core metabolism of *V. natriegens* in the glucose condition are shown. The majority of the k_app_ estimates converge within a narrow band, independent of their initial estimate. Similar trends hold true for the other conditions as well (Supplementary Figures S33 – S37). The median coefficient of variation of the k_app_ estimates across all the carbon conditions ranges from 6 % to 19 %. Across all the metabolic modules in Figure 6, the k_app_ estimates both increase or decrease to within their final range, suggesting that the parameterization scheme is not biased in a particular direction. The primary reason some k_app_ values exhibit a wider range in their final estimates is due to measurement uncertainty, e.g. the uncertainty shown in Figure 3D translates to uncertainty in Figure 6D. In other cases structural artifacts in the model give rise to uncertainty, e.g. aconitate hydratase (ACONT) is composed of two reactions in the model, both catalyzed by the same enzyme (i.e. there are two k_app_’s for this single enzyme). Thus, not enough measurements are available to parameterise both k_app_ values independently. Despite this, the tight convergence seen in Figure 6 strongly suggests that there is a small basin of attraction of kinetic parameters that minimize the total loss of Equation (7).

**Figure 6:**
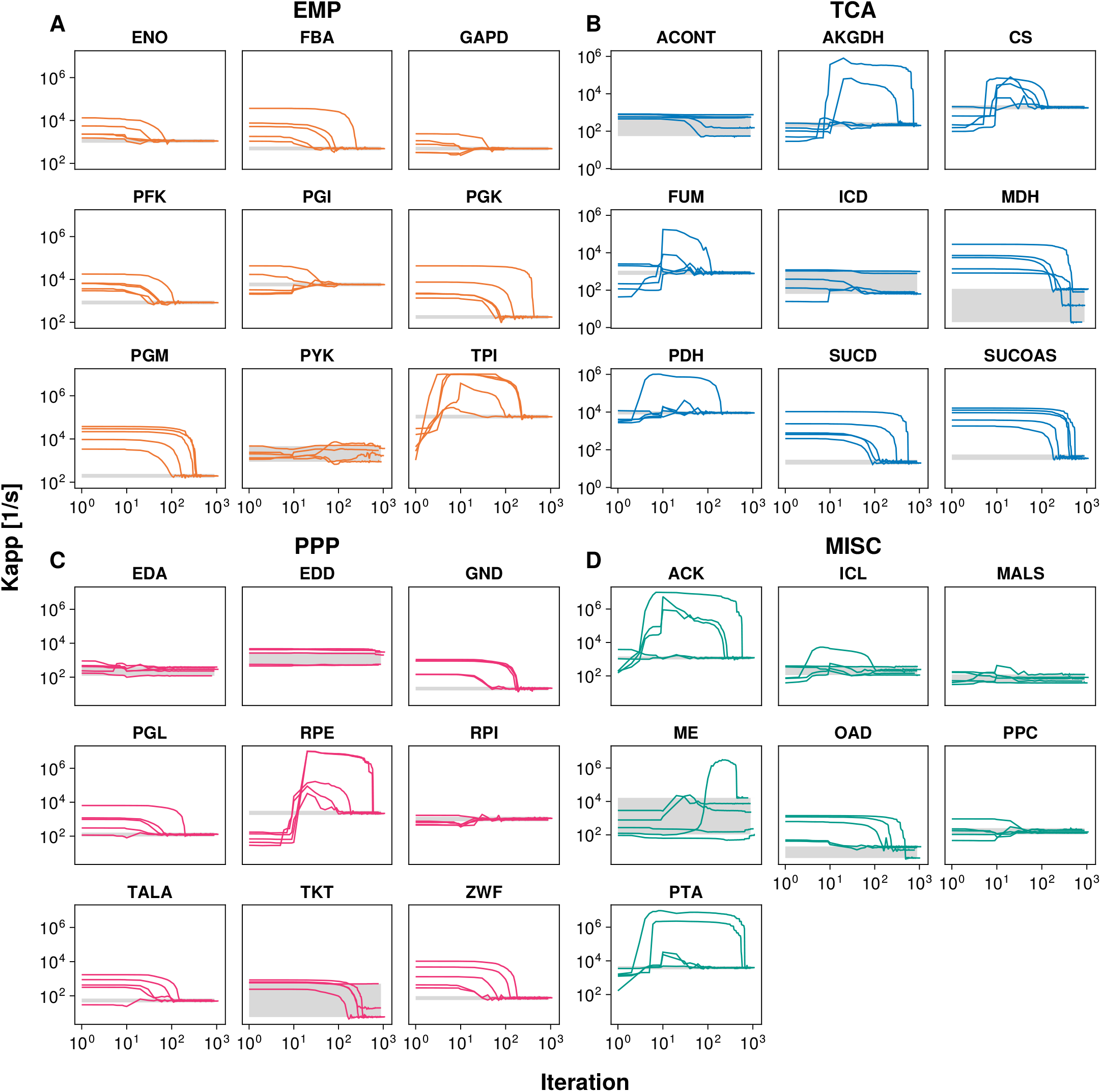
Kinetic parameter estimates converge to narrow ranges independent of initial conditions. For **A - D** the k_app_ estimates over optimization iterations are plotted as lines, and the extrema of the final estimates are plotted as a grey bands. The glucose condition is used here, see Supplementary Figures S33 – S37 for the other conditions. The k_app_ estimates in the Embden-Meyerhof-Parnas pathway (EMP, glycolysis) (**A**), citric acid cycle pathway (TCA) (**B**), pentose phosphate pathway (PPP) (**C**), and miscellaneous metabolic modules (MISC: anaplerotic, glyoxylate shunt, and acetate overflow) (**D**) largely converge within 1000 iterations of the fitting optimization problem (Equation 7). Exceptions are largely localized to reactions with high uncertainty, as shown in Figure 3. ACONT: Aconitate hydratase, ACK: Acetate kinase, AKGDH: 2-Oxogluterate dehydrogenase, CS: Citrate synthase, EDA: 2-dehydro-3-deoxy-phosphogluconate aldolase, EDD: 6-phosphogluconate dehydratase, ENO: Enolase, FBA: Fructose-bisphosphate aldolase, FUM: Fumarase, MDH: Malate dehydrogenase, GAPD: Glyceraldehyde-3-phosphate dehydrogenase, GND: Phosphogluconate dehydrogenase, ICD: Isocitrate dehydrogenase, ICL: Isocitrate lyase, MALS: Malate synthase, ME: Malic enzyme, OAD: Oxaloacetate decarboxylase, PDH: Pyruvate dehydrogenase, PGK: Phosphoglycerate kinase, PFK: Phosphofructokinase, PGI: Glucose-6-phosphate isomerase, PGL: 6-phosphogluconolactonase, PGM: Phosphoglycerate mutase, PPC: Phosphoenolpyruvate carboxylase, PTA: Phosphotransacetylase, PYK: Pyruvate kinase, RPE: Ribulose 5-phosphate 3-epimerase, RPI: Ribose-5-phosphate isomerase, SUCD: Succinate dehydrogenase, SUCOAS: Succinyl-CoA synthetase, TALA: Transaldolase, TKT: Transketolase, TPI: Triose-phosphate isomerase, ZWF: Glucose 6-phosphate dehydrogenase.

### Multi-omic data informed parameter estimates improve model predictions

Subsequent to the parameter estimation procedure, the final condition specific k_app_ estimates per reaction is the mean over the five replicates for the associated condition (Supplementary Data 6). Thus, each condition has a final k_app_ parameterisation, and using Equation (8) we can find the k_max_ of each reaction across all the conditions.

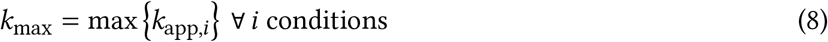

For the final condition dependent k_app_ datasets, we distinguish between their predictive and explanatory power. We measure the predictive power of a parameterisation as the loss between flux and protein predictions relative to measurements when solving Equation (2). This is the classical growth maximizing enzyme constrained metabolic model, whose objective is different to that of the inner problem in Equation (7). This definition reflects a typical use-case of enzyme constrained models when intracellular measured data are not available. In contrast, we measure the explanatory power of the parameterisation as the solution of the inner problem in Equation (7) given a parameterisation. In the context of our work, this best reflects the underlying biology as captured by the model, since the optimization problem seeks to minimize the difference between measurements and predictions. The only difference between these two problems is the objective function. The former maximizes growth, while the latter minimizes the difference between predictions and measurements, i.e. given a kinetic parameterisation, how close can the predictions of the model get to the observations.

The biological explanatory power of the parameter estimation model was used to investigate the enzyme kinetics of *V. natriegens* further. The distribution of the final k_app_ estimates for each condition is dependent on the growth rate of the condition. As expected, the faster growing conditions have elevated kinetics with the median k_app_ per condition ranging from 89 s^−1^ to 180 s^−1^ (Figure 7A). On a matched enzyme basis, the k_max_ estimates for *V. natriegens* are in median 14-fold faster than those of *E. coli* [10]. The median k_max_ for *V. natriegens* and *E. coli* is 210 s^−1^ and 16 s^−1^. The interquartile range of the k_max_ values for *V. natriegens* is 33 s^−1^ to 763 s^−1^, and for *E. coli* 6 s^−1^ to 43 s^−1^. These trends extend across every metabolic module contained in the model, demonstrating that *V. natriegens* has signficantly faster metabolic enzymes (Figure 7B).

**Figure 7:**
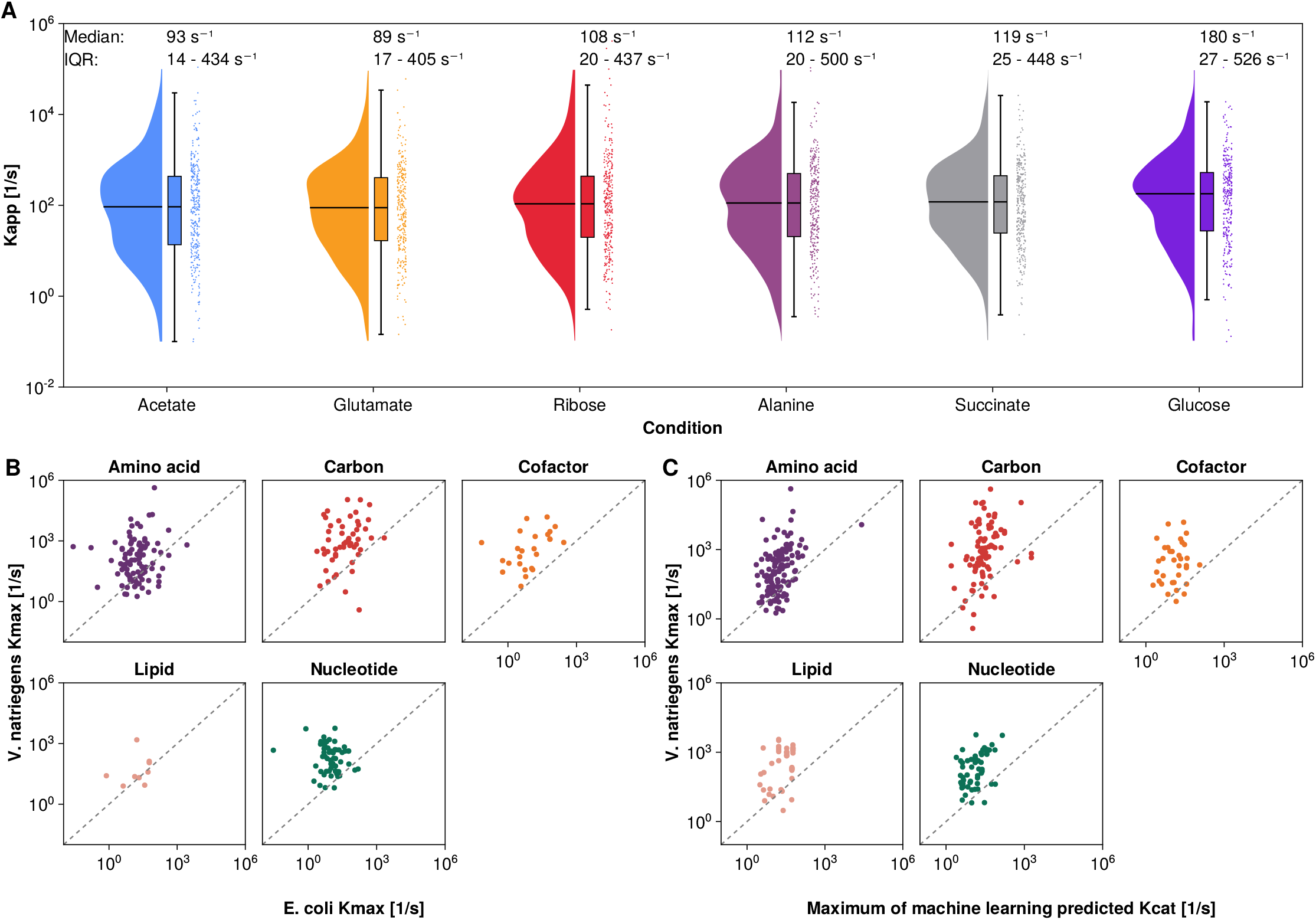
Distribution of optimal kinetic parameter estimates. The average value for each five replicates with the lowest total loss was taken as the final estimate for the k_app_ per reaction. The maximum apparent turnover number (k_max_) was estimated using Equation 8. **A:** The k_app_ estimates across each condition reveal that faster growth conditions lead to faster enzyme kinetics. Conditions ordered in increasing growth rate, from acetate (0.63 ± 0.01 h^−1^) to glucose (1.74 ± 0.04 h^−1^). **B:** Comparing the k_max_ between *V. natriegens* and *E. coli* reveals that the metabolic enzymes of *V. natriegens* are 14-fold faster in median, explaining its faster growth rate. **C:** Machine learning parameter estimates were unable to accurately predict the enhanced kinetics of *V. natriegens*, consistently underestimating its kinetics. The maximum k_cat_ estimate per reaction of two state-of-the-art machine learning models was used as the reference [29, 28]. IQR: interquartile range.

Comparing the k_max_ estimates derived here to k_cat_ predictions made by two state-of-the-art machine learning models reveals stark differences (Figure 7C). A consensus machine learning k_cat_ estimate was used in this work, which is the maximum k_cat_ prediction from two models that make use of protein sequence and reaction metabolite information to generate kinetic predictions [29, 28]. The median k_cat_ estimate of the machine learning models is 18 s^−1^, and the interquartile range is 10 s^−1^ to 34 s^−1^. These predictions align poorly with the measured data presented here, including the k_app_ estimates of monomeric enzymes where both the flux and protein concentration was directly measured. In contrast, the predictions align much better with the kinetic estimates for *E. coli*, likely reflecting that the training data was understandably dominated by *E. coli* measurements.

Using the optimized k_app_ parameters, the flux predictions of the context specific models were investigated and compared to measurements (Figure 8). As explained, we dilineate between the predictive and the explanatory power of the models by using a different objective for Equation (2) in each case. In the predictive case, the objective is growth maximization (*µ*), and in the explanatory case the objective is the total loss, as described by Equation (6). Here, context specific models mean that the models only use reactions for which k_app_ estimates are available after paramater optimization using Equation (7) (in short, metabolically active reactions in the respective carbon source, see the material and methods section for more details on how they were constructed). We purposefully do not use the k_max_ estimates here, as those estimates ignore the effect of metabolic regulation on the flux profile of *V. natriegens* when growing on a specific carbon source, which is large, as discussed later. The explanatory model is much better able to predict flux measurements (median relative error is approximately 3 % of the measured flux), which is expected since it is the inner problem of the parameter estimation model (left hand side of Figure 8). By extending the analysis to the growth maximization objective to test the predictive accuracy of the model, we find that the errors increase (median relative error of 44 %). This is due to a technical artifact of enzyme constrained flux balance analysis. Briefly, an optimal solution for an enzyme constrained growth maximization model can have at most as many active elementary flux modes (EFMs) as capacity constraints [30, 31]. Inspecting the intracellular flux profiles of, e.g. the glucose condition, we notice that more than two EFMs must be active (besides acetate fermentation, both the EMP and ED pathway are active). Thus, when optimizing for growth maximization the model shifts the enzyme capacity to at most two EFMs. Since we only have a total- and membrane capacity bound in Equation (2), this causes incorrect predictions as certain reactions are switched off freeing up enzyme capacity for other pathways. This is most prominent in the large relative errors associated with the ED pathway (EDA: 2-dehydro-3-deoxy-phosphogluconate aldolase, EDD: 6-phosphogluconate dehydratase) on the right hand side of Figure 8. Despite this technical limitation of enzyme constrained flux balance analysis, the growth maximization models accurately predict the fluxes near the entry point of their respective carbon source. For example, the glucose condition has the lowest relative errors in glycolysis (median 2 % in the EMP pathway vs. 75 % in the PPP and TCA pathways), the ribose condition in the PPP system (median 8 % in the PPP system vs. 17 % in the EMP and TCA pathways), and the gluconeogenic substrates in the TCA pathway. There is also a modest trend for faster growing conditions to have less overall relative error, likely because the parsimonious enzyme usage assumption embedded in enzyme constrained flux balance analysis models holds better.

**Figure 8:**
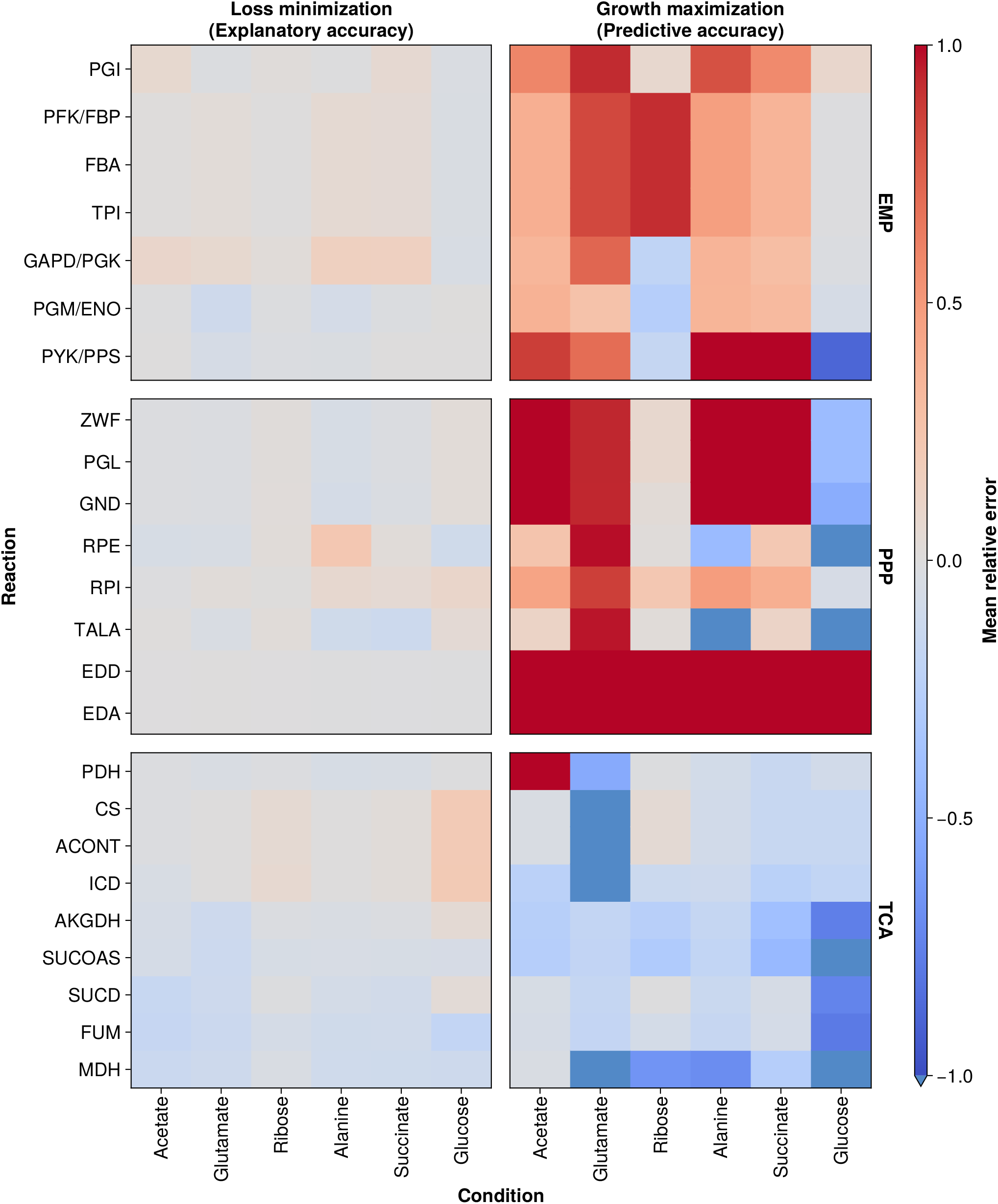
The predictive power of the condition specific kinetic parameters is highest in metabolic modules closer to the associated carbon source. Here the predictions of two models are shown: “Loss minimization” corresponds to the predictions of the inner enzyme constrained flux balance analysis model in Equation 7, using the final k_app_ estimates of each condition, while “Growth maximization” corresponds to the predictions using Equation 2 with the same k_app_ estimates. Reaction fluxes are also grouped by metabolic module. Mean relative error is calculated by 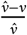 for a predicted (*𝓋*) and measured (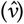) flux. ACONT: Aconitate hydratase, AKGDH: 2-Oxogluterate dehydrogenase, CS: Citrate synthase, EDA: 2-dehydro-3-deoxy-phosphogluconate aldolase, EDD: 6-phosphogluconate dehydratase, FBA: Fructose-bisphosphate aldolase, FUM: Fumarase, MDH: Malate dehydrogenase, GAPD/PGK: Glyceraldehyde-3-phosphate dehydrogenase/Phosphoglycerate kinase GND: Phosphogluconate dehydrogenase, ICD: Isocitrate dehydrogenase, ICL: Isocitrate lyase, MALS: Malate synthase, ME: Malic enzyme, OAD: Oxaloacetate decarboxylase, PDH: Pyruvate dehydrogenase, PFK/FBP: Phosphofructokinase/Fructose-bisphosphatase, PGI: Glucose-6-phosphate isomerase, PGL: 6-phosphogluconolactonase, PGM/ENO: Phosphoglycerate mutase/Enolase, PPC/PPCK: Phosphoenolpyruvate carboxylase/Phospho-enolpyruvate carboxykinase, PTA/ACK: Phosphotransacetylase/Acetate kinase, PYK/PPS: Pyruvate kinase/Phosphoenolpyruvate synthase, RPE: Ribulose 5-phosphate 3-epimerase, RPI: Ribose-5-phosphate isomerase, SUCD: Succinate dehydrogenase, SUCOAS: Succinyl-CoA synthetase, TALA: Transaldolase, TKT: Transketolase, TPI: Triose-phosphate isomerase, ZWF: Glucose 6-phosphate dehydrogenase.

Finally, focussing only on the predictive case using Equation (2), we investigated the effect of different kinetic paremerizations on the phenotypic predictions of the context specific models. Using the machine learning predicted kinetic constants (k_cat_ values) proved to be challenging, as they typically led to infeasible solutions given the capacity bounds of the model. These capacity bounds are in general growth limiting, but a minimum maintenance energy (ATPM) is also required by the model; on average, across all conditions, the ATPM reaction was 97.5 mmol g_DW_^−1^. This latter bound could not be overcome using the machine learning predicted k_cat_ values. In contrast, the multi-omic based k_app_ and k_max_ estimates could accurately recapitulate the measured growth rate (Figure 9A). For these simulations the extracellular carbon uptake rate was constrained to its measured value, because transporter kinetics were excluded from the parameterization simulations. Consequently, the use of the k_max_ values as kinetic constants did not materially affect the growth rate. However, significantly less protein was required to fully utilize the carbon uptake for growth (Figure 9B). This is due to the large range of k_app_ values (on average spanning one order of magnitude) across conditions (Figure 9C). Enhanced enzyme kinetics equates to more flux per enzyme, and the substrate uptake bound is hit before the enzyme capacity bounds. This issue is significantly attenuated when using (context specific) k_app_ estimates, with both the membrane and total capacity being nearly saturated (the total capacity is on average 92 % saturated in Figure 9B). Using the k_app_ estimates, the protein capacities are only modestly subsaturated due to the EFM protein re-allocation artifact discussed previously. Interestingly, the combination of Figures 8 and 9 imply that despite the fast growth of *V. natriegens*, the organism does not allocate its proteome for maximum growth rate (assuming the accuracy of the models). Despite this technical limitation, the context specific parameterisation accurately predicts the growth rate.

**Figure 9:**
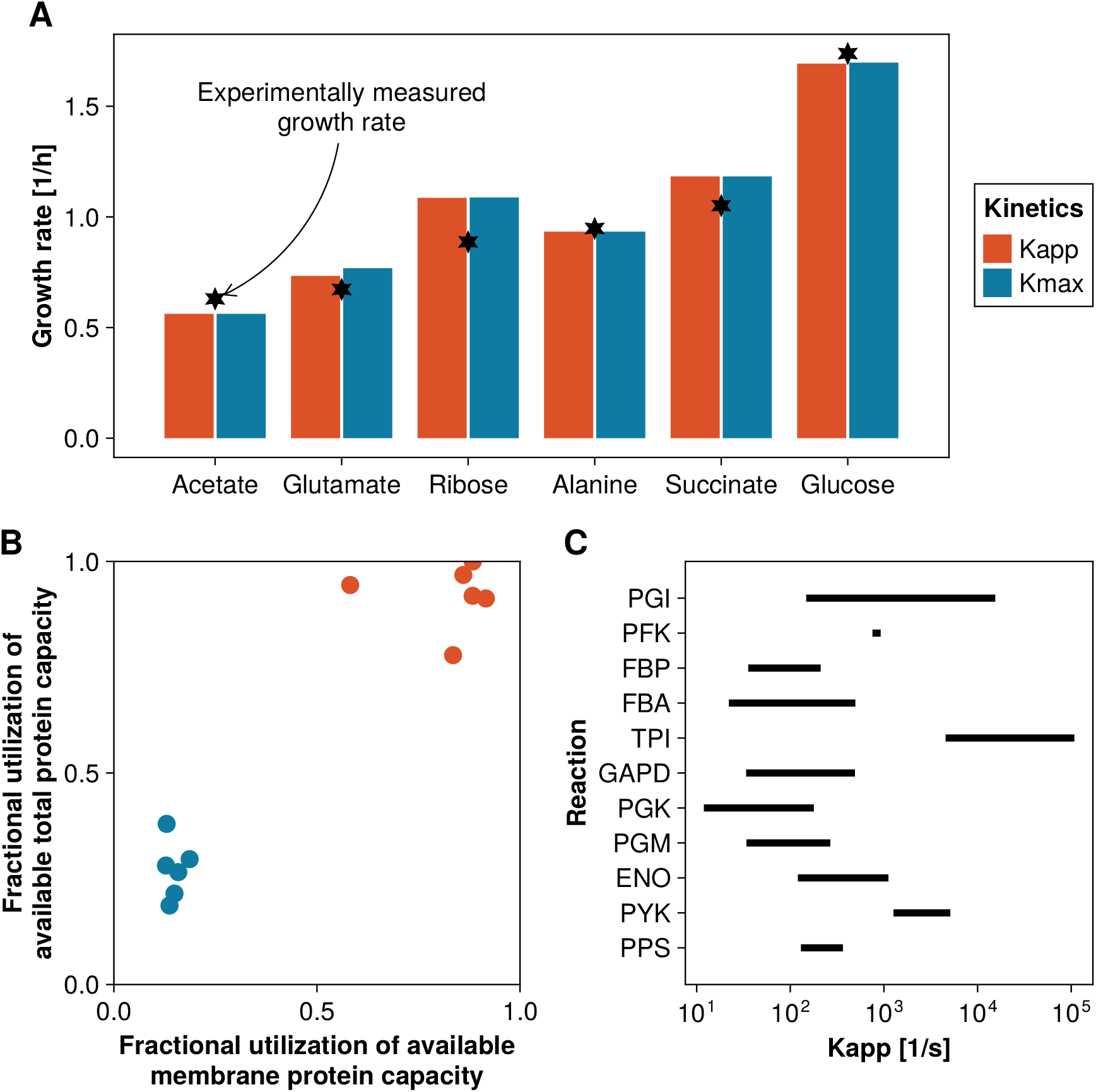
Context specific models with multi-omic informed kinetic parameters accurately predict growth rate measurements. The growth rate of *V. natriegens* was predicted using different kinetic parameterisation schemes: k_app_ corresponds to the optimized parameters for each condition, while k_max_ is the maximum k_app_ across all conditions per reaction. In all cases Equation (2) was solved with the corresponding context specific model. The extracellular uptake flux of carbon was constrained to the measured value. **A:** Growth rate predictions align very well to experimental measurements. **B:** Corresponding to the simulation of **A**, the utilization of protein capacity in the membrane and total pool was recorded. Using the k_max_ kinetic estimates, significant excess protein is available, but since the extracellular uptake flux is limited, this does not lead to an increased growth rate. **C:** The range of k_app_ estimates in glycolysis typically span an order of magnitude, suggesting significant substrate specific metabolic regulation. Similar results hold across other metabolic modules. ENO: Enolase, FBA: Fructose-bisphosphate aldolase, FBP: Fructose-bisphosphatase, GAPD: Glyceraldehyde-3-phosphate dehydrogenase, PGK: Phosphoglycerate kinase, PFK: Phosphofructokinase, PGI: Glucose-6-phosphate isomerase, PGM: Phosphoglycerate mutase, PPS: Phosphoenolpyruvate synthase, PYK: Pyruvate kinase, TPI: Triose-phosphate isomerase.

In sum, these results highlight the benefit of using omic-based measurements to parameterize models. The multi-omic data fusion approach used here successfully improves the predictive accuracy of metabolic models, and demonstrates a systematic approach to extend the kinetic estimates of microbes using increasingly abundant omic data.

## 3 Discussion

Combining ^13^C MFA estimated intracellular fluxes and quantitative proteomics for *V. natriegens* grown under a variety of culture conditions enables the direct calculation of *in vivo* apparent enzyme turnover numbers. The advantage of this approach compared to classic *in vitro* assays includes simpler scalability and naturally incorporating the effect of cellular context on turnover number estimates. Indeed, as predicted by Davidi et al. [9] 10 years ago, the increasing robustness of quantitative proteomics and intracellular flux estimates has led to k_app_ estimates for *E. coli* [10] and *Saccharomyces cerevisiae* [32], which in both cases led to improvements in metabolic modeling predictions.

Here we present data that largely affirms the predictions of modern microbial growth laws: the ribosomal proteomic fraction of cellular dry mass increases linearly with growth rate, the total protein density (capacity) is independent of growth rate, and the combination of these two constraints effects a trade-off in enzyme abundance (anabolic vs. catabolic) with growth rate (Figure 2). Recapitulating previous work, we find that the coarse proteome allocation does not differ widely between *E. coli* and *V. natriegens*, but specific culture conditions cause significant regulation at the pathway level [7]. We show that *V. natriegens* is capable of channeling very high fluxes through its metabolism, but that its intracellular flux profile resembles that of *E. coli*, which is not surprising given their similar metabolism (Figure 3). Integrating the proteomic and flux measurements yields *in vivo* k_app_ estimates, of which the median k_max_ is 14-fold faster than *E. coli* on a matched-enzyme basis (Figure 7). These data supply further evidence that *all* its enzymes, including its ribosomes, are faster than those of other slower growing microbes. Given these findings, it is tempting to speculate that *Geobacillus* LC300, an even faster growing bacterium, may have enzymes that are kinetically faster than those of *V. natriegens*.

This naturally raises several questions: (1) why do all organisms not evolve faster enzymes? We cannot offer conclusive evidence here, but it is probable that enzyme kinetics (and species growth rate) are niche adapted. It may just not make evolutionary sense for *E. coli* to grow as fast as *V. natriegens*, as overgrowth in its natural microbiome (the human gut) would likely hurt the fitness of its host or other symbiotic microbes, thereby creating a natural dampening effect on its enzyme evolution. (2) What is the mechanistic cause of the faster enzyme kinetics in *V. natriegens*? Previous data for *E. coli* has shown that enzymes tend to be substrate saturated [33], suggesting that either thermodynamic or structural factors are more likely than substrate saturation effects to explain the higher k_app_ values. In the case of thermodynamics, most enzymes operate far from their equilibrium, reducing the impact of thermodynamic driving force on flux. The Haldane relationship suggests that there is a trade-off between reaction speed and affinity, and recent work has found that some enzymes (e.g. RuBisCO) sacrifice speed for specificity [34]. It may be that *V. natriegens* sacrifices affinity/specificity for speed, but further intracellular metabolomic measurements would be necessary to adjudicate this explanation. Alternatively, aligning the metabolic enzymes of *E. coli* and *V. natriegens* reveals substantial dissimilarity at the amino acid level. It seems likely that structural effects might be more important than saturation or thermodynamics in explaining the kinetic difference between the enzymes of the two microbes. In addition, allosteric regulation may also play a role in the kinetic differences. (3) Would heterologous expression of *V. natriegens* enzymes in e.g. *E. coli* lead to higher fluxes in the mutant strain? It seems unlikely that this would occur for single enzyme replacements, as enzymes work in concert with each other. Thus, replacing e.g. *E. coli*’s pyruvate kinase with that of *V. natriegens* would likely not cause a measurable effect in its glycolytic flux. However, replacing an entire pathway might lead to significant metabolic adjustment.

The parameter estimation approach employed here is not limited to *V. natriegens*, and could easily be extended to any microbe. However, there are some caveats with this approach. Reliable proteomic and intracellular flux measurements require careful experimental design and advanced evaluation to ensure their fidelity. In particular, for the ^13^C MFA flux estimation method used here, the organism must grow stably enough to reach metabolic and isotopic steady state. Alternatively, non-stationary ^13^C MFA can be used [35], which also becomes necessary in situations such as C_1_-based carbon sources [36]. Further, flux imputation directly depends on the accuracy of the genome-scale metabolic model being used. This is a particular concern given the ubiquity of automated model construction tools that produce models invariably requiring further curation, which is usually only sparingly performed. Moreover, the accuracy of k_app_ predictions is also directly connected with the fidelity of the gene reaction rules, and are centrally important in fusing the proteomics data to fluxes.

Despite these caveats, integrating heterogenous omics data with enzyme-constrained flux modeling unlocks a wealth of information that can be used to interrogate the metabolism of microbes in diverse growth conditions — furthering our understanding of their evolutionary adaptations. While machine learning tools are training data limited, we expect their predictions to lag behind measurement schemes like the one used here. This increases the importance of basing model predictions on measured data as much as possible, as it reduces the need for *ad hoc* modeling assumptions, increasing the generalizability and applicability of enzyme constrained metabolic models to bioengineering endeavors. In sum, we demonstrate the increasing importance of using quantitative systems biology to investigate microbial metabolism by *inter alia* affirming the mechanism *V. natriegens* uses to accelerate its growth and measuring hundreds of enzyme turnover numbers simultaneously.

## 4 Materials and Methods

### Strain, media, and culture conditions

Wildtype *V. natriegens* (ATCC 14048, DSM 759) was used in all experiments. *V. natriegens* was stored at −80 °C in 25 % (vol/vol) glycerol stocks, and streaked out on LBv2 medium plates supplemented with 18 g L^−1^ agar prior to experiments. LBv2 was made as previously described [37]. Unless otherwise indicated, VN medium [2] was used for all experiments, albeit with a reduced 3-(N-morpholino)propanesulfonic acid (MOPS) concentration of 10 g L^−1^ and different carbon sources as listed below. Only for the ^13^C MFA were isotopically enriched carbon sources used: D-Glucose ([1,2]-^13^C_2_), D-Ribose ([5]-^13^C), L-Alanine ([1]-^13^C), Glycerol ([1,3]-^13^C_2_), L-Glutamic acid ([1,2]-^13^C_2_), Succinic acid ([1,4]-^13^C_2_), and Acetic acid ([1]-^13^C). The ^13^C tracers were purchased from Cambridge Isotope Laboratories (USA), with chemical purity specification for each carbon source ≥ 98% and the isotopic enrichment of each carbon source ≥ 99%. Otherwise, ^12^C carbon sources were purchased from Sigma Aldrich. A culturing strategy, detailed in Supplementary Sections S2.1 and S3.1, was followed to ensure the cells were in balanced, exponential growth at the time of harvesting for all experiments. Biological replicates were made in triplicate, except for the proteomic experiments, which made use of quadruplicates. Experiments were conducted either in a BioLector Pro (Beckman Coulter), using a 48 well flower plate (M2P-MTP-48-B, Beckman Coulter), sealed with a gas permeable foil (M2P-F-GPR48-10, Beckman Coulter) or in shaking glass flasks as detailed below. For each replicate, the corresponding wells (flasks) were inoculated from a well mixed 50 mL seed culture in fresh medium maintained at 37 °C in exponential growth for at least 10 doublings, ensuring that each well (flask) had the same starting OD_600_ and the culture conditions did not change prior to inoculation. The BioLector chamber was humidity and temperature (37 °C) controlled, with an agitation rate of 1000 rpm to enable maximum oxygen transfer to *V. natriegens*. Flasks were shaken at 230 rpm on an orbital shaker. For the BioLector plates, dissolved oxygen and pH were monitored using the integrated optodes, to ensure *inter alia* that oxygen was not limiting during any of the experiments. Proteomic and ^13^C MFA metabolomic samples were all harvested during mid-exponential growth, at an OD_600_ of between 0.5 – 0.8. We found this to be important as the rapid growth of *V. natriegens* tended to deplete the oxygen concentration in the medium quickly near stationary phase. Extracellular metabolomic samples were harvested at regular intervals during a growth curve to measure extracellular fluxes.

### Proteomics sample preparation

Samples were cultured as described, and harvested in quadruplicate (see Supplementary Section S2.1 for details). Thereafter, they were washed twice in TBS (7 min at 4500 g, 4 °C), and the pellet resuspended in 150 mM Tris, 50 mM DTT and 0.1 % for lysis. The samples were incubated at 4 °C for 10 min on a rotary shaker, before being frozen overnight at −20 °C. The next day an additional lysing step was performed by ultrasonication (3 min in 6 cycles of 30 s ultrasonication and 30 s break) and frozen at −80 °C for later processing. For LC-MS analysis a modified magnetic bead-based sample preparation protocol was used as described in [38]. Briefly, 10 µg *V. natriegens* protein per sample was reduced by adding 5 µL 100 mM DTT (dithiothreitol) and shaken for 20 min at 56 °C and 1000 rpm, followed by alkylation with the addition of 6.67 µL 300 mM IAA and incubation for 15 min in the dark. A 20 g L^−1^ bead stock of 1:1 Sera-Mag SpeedBeads was freshly prepared and 5 µL were added to each sample. Afterwards, 42 µL ethanol was added and incubated for 15 min at 24 °C. After three rinsing steps with 80 % ethanol and one rinsing step with 100 % acetonitrile, beads were resuspended in 50 mM TEAB buffer and digested with final 1:50 trypsin at 37 °C and 1000 rpm overnight. Additional digestion was carried out by adding trypsin (final ratio 1:50) and shaking at 37 °C and 1000 rpm for 4 h. The supernatants were collected and 500 ng of each sample digest were subjected to LC-MS.

### Proteomics LC-MS/MS analysis

For the LC-MS acquisition an Orbitrap Fusion Lumos Tribrid Mass Spectrometer (Thermo Fisher Scientific) equipped with a FAIMS module coupled to an Ultimate 3000 Rapid Separation liquid chromatography system (Thermo Fisher Scientific, Idstein, Germany) equipped with an Acclaim PepMap 100 C18 column (75 µm inner diameter, 25 cm length, 2 µm particle size from Thermo Fisher Scientific) as separation column and an Acclaim PepMap 100 C18 column (75 µm inner diameter, 2 cm length, 3 µm particle size from Thermo Fisher Scientific) as trap column was used. A LC-gradient of 180 min was applied. The FAIMS module was operated with a carrier gas flow of 4.5 L/min and the compensation voltage (CVs) was set to −50 V. MS survey scans were performed in the Orbitrap analyzer at a resolution of 60,000 and target values of 400,000 ions and a maximum fill time of 100 ms over a mass range from 380 to 985 m/z. For DIA MS/MS scans the resolution was set to 15,000. Sixty sequential precursor isolation windows, each with a window size of 11 m/z, were used to cover the MS1 scanning range.

### Proteomics data analysis

Data analysis was performed using Spectronaut (version 19.8, Biognosys, Schlieren, Switzerland). All RAW files were processed in directDIA mode using the default Spectronaut workflow parameters. MS/MS spectra were searched against a *V. natriegens* protein sequence database generated from the predicted proteome of the NCBI genome assembly ASM145625v1 (RefSeq annotation, accessed in 2025). Enzyme specificity was set to trypsin/P, allowing up to two missed cleavages. Carbamidomethylation of cysteine residues was defined as a fixed modification, while methionine oxidation was included as a variable modification. Standard Spectronaut settings for precursor and fragment ion mass tolerances were applied automatically by the software. Peptide and protein identification confidence was controlled using the built-in target–decoy approach implemented in Spectronaut. False discovery rates (FDRs) were controlled at 1 % at both the precursor and protein levels. Protein inference and quantification were performed using the default Spectronaut directDIA algorithms and normalization procedures. Unless otherwise stated, all other analysis parameters were kept at their default values. Briefly, inconsistent and interfering peptide signals were reduced across runs by applying the default q-value threshold filter implemented in Spectronaut. Major group quantities were determined using the Mean Peptide Quantity approach combined with Top N peptide selection (maximum 3, automatically selected per protein). For each protein group, quantitative values were calculated as the mean abundance of the Top N most intense peptides passing the applied quality filters. Cross-run normalization was enabled using the automatic normalization settings. The default local normalization strategy based on local regression normalization was used [39]. The normalization set was selected automatically using precursor profiles ranked by completeness across runs. For more details regarding replicate handling and consistency checks see the Supplementary Section S2. The mass spectrometry proteomics data have been deposited to the ProteomeXchange Consortium via the PRIDE [40] partner repository with the dataset identifier PXD078550.

### ^13^C MFA sample preparation

*V. natriegens* was cultivated as described in Supplementary Section S4.1. At least six samples were harvested in an OD_600_ range of 0.5 – 0.8. The samples were washed three times (1 min at 4500 g, washing buffer), and the resultant pellets frozen at −80 °C for later batch processing. The overall workflow of protein hydrolysis and GC-MS measurement was adapted from [41]. A cell culture pellet equivalent of 0.3 OD_600_ units was washed 3 times with deionized water and dried over night in a vacuum centrifuge. Subsequently, 500 µL 6 M HCl was added to the dried material, and the mixture was incubated for 24 h at 105 °C to completely hydrolyze the biomass proteins. After that, the samples were dried under a nitrogen stream. The free amino acids were redissolved in 150 µL 0.1 % formic acid. Thereafter, 15 µL of sample was then mixed with 50 µL of methanol and dried again in glass inlets in a vacuum centrifuge. The glass inlets were transferred to GC-MS glass vials and placed in a cooled autosampler (MPS, Gerstel) at 4 °C. Prior to measurement the samples were automatically derivatized by adding 50 µL of acetonitrile and 50 µL N-(tert-butyldimethylsilyl)-N-methyl-trifluoroacetamide (TBDMS, Sigma-Aldrich) at 30 °C for 70 min under agitation. The resulting TBDMS derivatives were used in following GC-MS analysis.

### ^13^C GC-MS based isotopologue profiling of proteinogenic amino acids

The resulting TBDMS derivatives were analyzed with GC-MS according to [42] with some modifications. Briefly, 1 µL of derivatized compounds was injected with an automatic liner exchange system in conjunction with a cold injection system (Gerstel) (ramping from 50 °C to 250 °C at 12 °C s^−1^) into the GC with a helium flow of 1 mL/min. Chromatographic separation was performed using a 5977B GC/MSD system (Agilent Technologies) with a HP-5MS column with 5 % phenyl methyl siloxane film (Agilent 19091S-433, 30 m length, 0.25 mm internal diameter, 0.25 µm film). The oven temperature was held constant at 70 °C for 2 min and then ramped at 12.5 °C/min to 320 °C at which it was held constant for 5 min; resulting in a total run time of 27 min. Metabolites were ionized with an electron impact source at 70 eV and 200 °C source temperature and recorded in a mass range of m/z 60 to m/z 800 at 20 scans per second. Owing to degradation by acid hydrolysis, the amino acids tryptophan, arginine and cysteine could not be analyzed. Furthermore, acid hydrolyzation converted glutamine and asparagine to glutamate and aspartate. Therefore, results for aspartate and glutamate correspond to asparagine/aspartate and glutamine/glutamate, respectively. Metabolites were identified via MassHunter Qualitative (version b08.00, Agilent Technologies) by comparison of spectra to the NIST14 Mass Spectral Library (version 1a-v14). Peaks were integrated using MassHunter Quantitative (version b08.00, Agilent Technologies). Molar composition for each metabolite was calculated based on the contribution of each individual isotopologue in the analyzed fragments in Supplementary Table S1. The mass isotopomer/isotopologue data were corrected for naturally occurring stable isotopes by using uNAC (https://jugit.fz-juelich.de/IBG-1/ModSim/uNAC), standard deviations where calculated from corrected triplicates, with a lower bound on standard deviations of 0.004.

### ^13^C MFA modeling and simulation

A metabolic model of the central carbon metabolism of *V. natriegens* was formulated, comprising glycolysis, PPP, ED pathway, TCA cycle, glyoxylate shunt, ana- and cataplerotic reactions as well as amino acid synthesis and a cyclic exchange of CO_2_, partially based on the ^13^C MFA model by Long et al. [15]. The model was equipped with the necessary uptake and production reactions, accounting for all seven growth conditions (see Supplementary Figure S15). Cell growth was modeled as effluxes partitioned according the experimentally determined biomass composition [15], scaled with the observed growth rates. In total, the model contains 70 balanced and 8 extracellular metabolites: glucose, ribose, succinate, glutamate, alanine, glycerol, acetate, as well as CO_2_. The model contains 129 reactions of which 33 are bidirectional. The relatively high number of bidirectional reactions is explained by the unknown extent of the actual reversibility of the enzymatic reactions in the different cultivation conditions. From this model, seven condition-specific models were derived (of which six were fitted with sufficient quality). Reaction (bi)directionalities were constrained by credible prior knowledge, e.g. glutamate synthesis operates in the synthesis direction except for the case of glutamate-supplemented growth medium, where the reaction runs in reverse only. Extracellular rates as well as mass isotopomer data were incorporated in the models with their respective standard deviations. Additionally, wide upper and lower bounds to the fluxes were formulated to exclude implausible flux values. The models were formulated in the universal flux model language FluxML [43] in a single file, with one configuration for each isotope labeling experiment, where a configuration consists of reaction/flux constraints, the administered tracer and its purity, and the rate and mass isotopomer measurements. Model simulations were performed with the high-performance simulator 13CFLUX (version 3.0.0) [35]. To account for the high model flexibility due to the many unknown reaction bidirectionalities, and hence large number of expected non-identifiable exchange fluxes [44], a rigorous Bayesian multi-model approach was applied for flux estimation [45]. In short, Bayesian model averaging was applied to a model set with combinatorially many model variants, which allows marginalizing out exchange fluxes and generates robust net flux posterior probabilities. Technically, a specialized Markov chain Monte Carlo algorithm was used [46], implemented in the highly optimized polytope sampling toolbox hopsy [47]. For the net flux posterior distributions, mean marginal posterior values and 95 % credible intervals were derived. Predictive posterior checks were performed to assure that the model used for inference is consistent with the data.

### ^1^H-NMR for extracellular metabolomics

The ^1^H-NMR measurements of exometabolome samples were done as described previously with modifications [48]. In brief, a volume of 400 µL of the filtered sample was mixed with 200 µL of 0.2 M sodium hydrogen phosphate buffer solution, containing 30 % D_2_O (Euriso-Top) and 1.5 mM 3-trimethylsilyl-(2,2,3,3-D_4_) 1-propionic acid, sodium salt (TSP) (Carl Roth) in 5 mm NMR tubes (4”, Bruker Switzerland GmbH). The Bruker AVANCE-NEO 600 NMR spectrometer equipped with a SampleJet autosampler and a 5 mm QCI (H/C/N/F) cryo probe was operated by TOPSPIN 4.0.9 software (Bruker Biospin). Metabolites were identified and quantified using AMIX Viewer 4.0.2 software (Bruker Biospin). The signal of TSP was calibrated to 0.0 ppm for spectra alignment. Signals of metabolites were compared to spectra of pure compounds from an in-house library for identification. The ERETIC signal was generated within all spectra by using external calibration with the ERETIC quantification tool based on PULCON [49]. Absolute concentrations of metabolites were calculated from integrals of metabolite peaks and the integral of the ERETIC signal. For the acetate condition and glucose conditions, a BBO 600 BB/H&F/D Cryo Prodigy probe was used instead, with TOPSPIN software version 4.5.0. In this latter case, an internal TSP signal was used for absolute metabolite quantification.

### Metabolic model construction

A novel genome-scale metabolic model was reconstructed from genomic and experimental information gathered during the course of this work following an established protocol [50]. Reaction gene associations were manually assigned for all primary metabolic reactions using multiple annotation sources, as described in Supplementary Section S5. The ensuant model is subject to an expansive MEMOTE-like [51] test suite for quality control. The model is available at https://github.com/wilkengroup/VibrioNatriegens.

### Parameter estimation algorithm and simulation settings

For each carbon source a condition specific model was constructed by removing inactive transporters (based on the NMR measured secretome, media composition, and intracellular flux estimates) and the consequently blocked metabolic reactions were also removed. All gene reaction rules were removed from the transporters because they are very uncertain, and membrane proteins are challenging to accurately measure. Subsequently, an enzyme constrained model was constructed with a total protein capacity ranging between 0.14 g g_DW_^−1^ to 0.16 g g_DW_^−1^ depending on the condition specific proteomic data, reflecting the mass fraction of primary metabolic enzymes accounted for in the model. The membrane capacity limitation was set at a constant 12 % of the total protein capacity, based on proteomic data. The contextual models were further simplified by removing all inactive reactions, which were identified by running an enzyme constrained flux balance analysis simulation of the aforementioned model, where the objective was growth rate maximization, and the fluxes were temporarily constrained to be within 40 % of their measured value for a specific carbon source condition. The resultant condition specific models were used in further analyses. Each reaction, which may be catalyzed by multiple isozymes, was allowed only a single k_app_, as discussed in the main text. Subsequently, the k_app_ values were optimized using Equation (7). Since the inner problem can be efficiently differentiated [12], it is possible to use the L-BFGS algorithm to rapidly solve the parameterisation model [52, 53]. For each condition, 200 replicates were solved, and the 5 replicates with the lowest loss were retained for further analysis. The measurements used in each replicate were based on the mean protein concentration from the proteomics data, and a randomly drawn sample from the posterior flux distribution per replicate. An initial k_app_ parameterisation for a reaction was randomly drawn from a uniform distribution spanning an order of magnitude increase or decrease from a machine learning generated k_cat_ prediction. A parameterisation optimization run was stopped when the change in objective value between two subsequent iterations decreased below 10^−8^ in an absolute sense. The predictive accuracy model, with the growth maximising objective of Equation (2), was based on the aforementioned contextual model, with the added constraint of extracellular fluxes set to their measured values.

### Computing resources

Julia Language [54] packages DataFrames [55], GLM [56], JuMP [57], COBREXA [58] and Makie [59] were used to analyze and plot the data used in this manuscript. Metabolic flux analysis was performed with the high-performance simulator 13CFLUX (version 3.0.0) [35].

## 5 Data availability

The primary data generated and analysed (quantitative proteomics, ^13^C MFA model and inferred fluxes, turnover number estimates) in this work are included with the published article and its supplementary information files. The raw proteomic data is available in the PRIDE repository with the dataset identifier PXD078550. Miscellaneous data generated during and/or analysed in the current study are available from the corresponding author on reasonable request. Exemplary scripts to reproduce the primary results of this work, including the context specific models and turnover number estimates, are available at https://github.com/wilkengroup/vibrio-natriegens-apparent-turnover-numbers-example-simulations.

## 6 Acknowledgements

The authors would like to gratefully thank Sina Wilken, Stefan Robertz, and Maurice Mager for thoughtful discussions; Nina Schulten for exploratory experimental work; Sofie Rüffer, Stefana Dukic, Vincent von Häfen, and Flora Schlüter for help constructing the model visualizations. We are thankful for excellent technical support by Elisabeth Klemp, Katharina Hinsen, Maria Graf and Katrin Weber from the CEPLAS Plant Metabolism and Metabolomics Laboratory. Finally, the authors would like to thank Wolfgang Wiechert for his continuous support.

## 7 Author Contributions

**SEW:** Conceptualization, Methodology, Software, Formal analysis, Investigation, Resources, Data Curation, Writing - Original Draft, Visualization, Supervision, Project administration, Funding acquisition **MB:** Investigation, Formal analysis, Data Curation, Writing - Review & Editing **MK:** Writing - Review & Editing **ABG:** Formal Analysis, Investigation, Writing - Review & Editing **KM:** Formal analysis, Investigation, Writing - Review & Editing **PL:** Methodology, Writing - Review & Editing **ML:** Writing - Review & Editing **KS:** Writing - Review & Editing **AS:** Formal analysis, Investigation, Writing - Review & Editing **PW:** Methodology, Formal analysis, Investigation, Data Curation, Writing - Review & Editing **KN:** Formal analysis, Investigation, Writing - Review & Editing, Resources **IA:** Writing - Review & Editing **OE:** Writing - Review & Editing

